# Temporal control of sgRNA library activation unlocks large-scale *in vivo* CRISPR screens

**DOI:** 10.1101/2025.08.14.670372

**Authors:** Silvia Fenoglio, Yi Yu, James Tepper, Lauren Grove, Alborz Bejnood, Samuel R Meier, Ashley H Choi, Hsin-Jung Wu, Annabel Devault, Shangtao Liu, Binzhang Shen, Tenzing Khendu, Hannah Stowe, Esther CH Uijttewaal, Minjie Zhang, Brian B Haines, Erik Wilker, Alan Huang, Daniel Schramek, Ulrich Elling, Xuewen Pan, Jannik N Andersen, Teng Teng

## Abstract

Functional genomics screens have illuminated genetic dependencies in cancer, but conventional *in vitro* approaches fail to capture vulnerabilities shaped by the tumor microenvironment. Here, we implement CRISPR-StAR (Stochastic Activation by Recombination), a next-generation inducible CRISPR screening platform for large-scale *in vivo* applications. The system uses a dual lox-based recombination system to enable guide-level normalization and clonal knockout phenotyping. To analyze the rich (barcode-embedded) sequencing output, we developed UMIBB, a superior Bayesian statistical framework for quantifying gene-level dropout and enrichment compared to conventional software packages. Screening a 30,000-sgRNA library in A549 xenografts, followed by clone representation and dropout correlation analyses, showed high fidelity and reproducibility with dropout phenotypes resolvable using as few as 30 tumors for this size library. Validation across multiple tumor models demonstrated that a single tumor can provide reliable, functional annotation for ∼1,000 genes leveraging intra-tumor library controls for normalization. Comparing *in vivo* and *in vitro* screens revealed that a substantial subset of tumor suppressor genes exerts strong phenotypic effects only observable *in vivo*. For example, single-gene knockout and transcriptomic profiling confirmed that KMT2C and KMT2D have contrasting impacts on tumor growth - an insight that would have been overlooked in standard cell culture. Looking ahead, CRISPR-StAR screening, combined with our user-friendly analysis pipeline available on GitHub (R-package), offer an integrated framework for creating *in vivo* dependency maps that can complement existing *vitro* datasets like DepMap and Achilles. Critically, our approach reduces animal use by up to 7-fold compared to conventional *in vivo* dropout screens. This represents a significant ethical and methodological advancement - achieving genome-scale resolution with far fewer animals and greater reproducibility.

## INTRODUCTION

Pooled *in vitro* CRISPR/Cas9 screening approaches, exemplified by large-scale efforts such as the DepMap project^1^, have significantly advanced our understanding of genetic dependencies in human cancer. These methodologies have been instrumental in expanding key concepts such synthetic lethality, lineage dependency and oncogene addiction providing a rich catalog of novel targets for therapeutic intervention^1,2^. For example, early functional genomics studies revealed that MTAP-deleted cancer cells are uniquely sensitive to PRMT5 inhibition^3–5^, providing the rationale for the development of MTA-cooperative PRMT5 inhibitors such as AMG193^6^, MTRX1719^7^, and TNG462^8^. Likewise, loss-of-function screens have also identified synthetic lethality between *STAG2* and *STAG1*^9,10^, as well as between *PTEN* loss and inhibition of mTORC2 components^11^. These findings underscore the power of functional genomics to uncover novel context-specific vulnerabilities in cancer.

Building on this foundational work in RNAi and CRISPR-based genetic screens^1,12^, we aimed to address the limitations of *in vitro* screens by enabling robust knockout screens in the noisier and more complex context of *in vivo* tumor biology^13,14^. While *in vitro* systems have been invaluable, they often fail to recapitulate critical interactions with the tumor microenvironment, immune system, and nutrient gradients, which are factors that profoundly influence tumor progression and therapeutic response^13,15–17^. By studying these dynamics *in vivo*, we hope to uncover novel context-dependent vulnerabilities that are undetectable in conventional tissue culture systems.

Historically, *in vivo* genetic screens have faced several major challenges including: (i) random library representation due to low cellular engraftment efficiency^18^, (ii) variability in sub-clonal expansion resulting from intratumoral heterogeneity, and (iii) overrepresentation of outlier clones, all of which can confound data interpretation^18^. To address these limitations, we implemented the CRISPR-StAR (Stochastic Activation by Recombination) screening platform^14^. This technology builds on the now-common use of unique molecular identifiers (UMIs) as clonal barcodes^19–23^. UMIs improve signal detection by enabling the identification of outlier clones and providing robust statistics for each sgRNA. However, this approach alone fails to overcome bottlenecks from inconsistent tumor take-rates or variable clonal growth. The CRISPR-StAR platform resolves these issues by integrating an inducible Cre/Cas9 dual system with the UMI barcodes. This provides a dynamic internal control for each tumor-initiating clone, enabling clonal ‘lineage’ tracking^14^ that mitigates library representation bias and accounts for tumor heterogeneity.

Several computational tools, such as MAGeCK^24^, ScreenBEAM^25^, BAGEL^26^ and JACKS^27^, have been developed to analyze the pooled CRISPR screen data. However, all existing methods are based on modeling sgRNA abundance distribution to determine gene-level impact, thus these methods may not properly model the discrete clonal abundance and sporadic clonal representation data from the CRISPR-StAR platform. To effectively analyze the complex next-generation sequencing (NGS) data generated by CRISPR-StAR, we developed a dedicated computational pipeline capable of extracting meaningful sgRNA dropout signals from individual tumors. This UMIBB pipeline (available on Github) utilizes embedded barcodes to achieve high-resolution clonal analysis. By conducting *in vivo* drop-out screens in the A549 lung cancer xenograft model, followed by statistical analysis and computational down sampling, we demonstrate that 30,000 sgRNAs can be functionalized *in vivo* using tumors from just 30 mice. This represents a log-order reduction in animal usage relative to conventional approaches^13^ made possible by the unique ability to track clonal dynamics and incorporate internal normalization, setting a new benchmark for efficiency and reproducibility in *in vivo* genetic screening^14^. Of note, our new UMIBB pipeline enables versatile, barcode-based clonal analysis for a wide range of applications, from CRISPR-StAR screens to any other study utilizing unique molecular identifiers (UMIs) such as barcoding studies^28^ or organoid studies^29–31^.

Notably, by including a parallel *in vitro* comparator arm, we identified a subset of genes whose loss promotes tumor growth exclusively *in vivo*. This set was enriched for epigenetic regulators such as members of the COMPASS family and SWI/SNF chromatin remodeling complexes. Validation studies using single gene knockout demonstrated that loss of *KMT2C* markedly accelerated tumor growth *in vivo*, whereas *KMT2D* knockout had the opposite effect, slowing tumor growth of A549 lung cancer cells. Neither gene exhibited a phenotype under 2D culture conditions, highlighting the unique insights that can emerge from large-scale *in vivo* genetic screening.

Collectively, the inducible CRISPR-StAR platform, coupled with a novel bioinformatic framework, substantially enhances the fidelity, scalability and interpretability of *in vivo* functional genomics screens. By temporally controlling sgRNA library activation in established tumors, the CRISPR-StAR platform overcomes key bottlenecks of traditional *in vivo* knockout screening methodologies^13,14^ with the CSTAR-UMIBB analysis package being available for use by the broader research community.

## RESULTS

### CRISPR-StAR: A controlled dual Lox-based inducible system for expression of matched-pair sgRNAs harboring unique molecular barcodes for clonal tracking

To enable large-scale *in vivo* screening we used an inducible approach that activates library expression in established tumors called CRISPR-StAR^14^. In this method, the sgRNA library vector contains a stop cassette inserted within the TRACR sequence, flanked by two pairs of lox sites (LoxP and Lox5171), preventing the expression of functional sgRNAs until Cre recombinase is activated (Fig. 1A). These lox pairs are mutually exclusive and recombine with similar efficiency^14^. Upon the addition of 4-hydroxytamoxifen (4-OHT) to induce CreERT2 activity, two constructs are generated with equal frequency. The active sgRNA construct is generated when the stop cassette is excised, restoring the TRACR sequence and allowing expression of a functional sgRNA, leading to gene knockout (KO). The inactive construct is produced when the TRACR sequence is removed, resulting in a non-functional sequence, which serves as a normalization control for the active counterpart (Fig. 1A). The ratio between the active vs inactive counts per sgRNA (A/I ratio) informs on the phenotype upon target gene KO (i.e. depletion or enrichment) (Fig. 1B).

**Figure 1.**
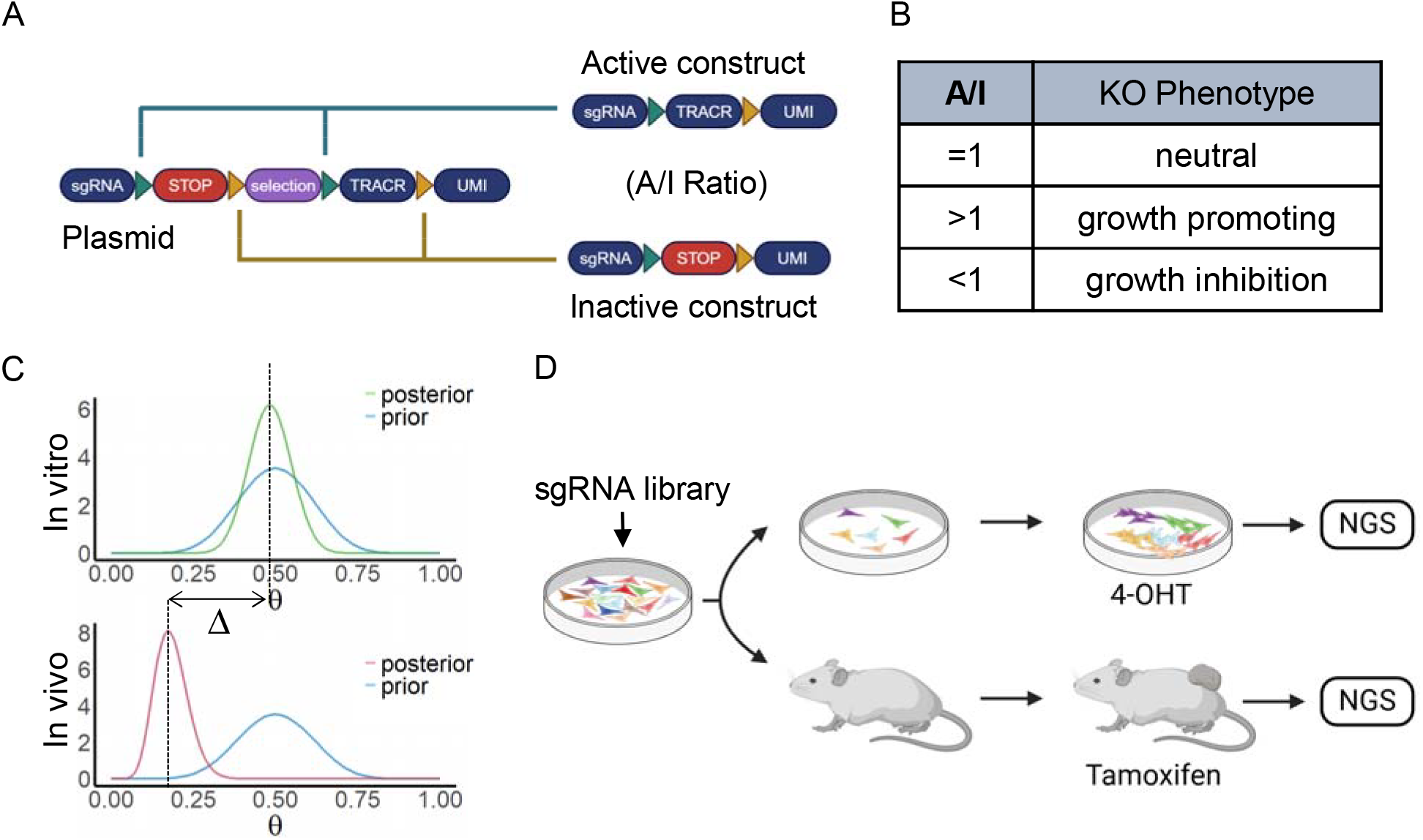
Overview of CRISPR-StAR system and the Bayesian analysis (UMIBB) of screening results. (A) CRISPR-StAR vector design enables Cre-mediated stochastic recombination via two pairs of incompatible Lox sites (LoxP and Lox5171), generating (with equal efficiency) either active (restored sgRNA-TRACR, resulting in gene knockout) or inactive (TRACR-deleted) constructs. (B) Knockout phenotypes are quantified as the ratio of active to inactive reads (A/I ratio). (C) Bayesian modeling framework for estimating wθ and Δ values in the UMIBB pipeline. (D) CRISPR-StAR screen setup. *In vitro*: 4-OHT (300 nM) was added to the culture media for 4 days to induce recombination. *In vivo*: Tamoxifen was administered to tumor-bearing mice to induce recombination.

By incorporating a barcode and implementing UMIs into each experimental vector, each clone is defined by the combination of its barcode and the sgRNA sequence in the endpoint dataset^20^. This design provides matched-pair controls, enabling clonal-level tracking of *in vivo* KO phenotypes. The paired control system ensures that any observed phenotypic differences are attributable to the specific gene knockout within the same micro-environment, improving the robustness and accuracy of the screening^14^.

### UMIBB: A Bayesian-based pipeline for clonal-resolved analysis of CRISPR-StAR datasets

To address the analytical challenges posed by the high complexity UMI data, we developed a computational pipeline, UMIBB (Bayesian Beta-binomial), designed to fully leverage the internal matched-pair controls and the wealth of independent clonal events captured in each tumor (Fig. 1B-C, Suppl Fig. 1). Since the ratio of active to inactive sgRNA counts (A/I ratio) serves as a direct measure of the sgRNA-induced phenotypes at the screening endpoint, we implemented a Bayesian inference model to robustly quantify dropout and enrichment phenotypes observed across clones. In this framework, each clone is treated as a Bernoulli trial, of which the probability of observing sgRNA depletion (A/I<1) or enrichment (A/I>1) can be modeled by a beta-binomial distribution (Fig. 1B-C). To summarize phenotypic trends across clones targeting the same gene, we defined a weighted-theta score (wθ) ranging from 0 to 1. This score reflects the weighted average frequency of enriched active constructs across all sgRNAs targeting a gene, with weights proportional to the number of informative clones per sgRNAs. A wθ score < 0.5 indicates a depletion phenotype, while scores > 0.5 indicate enrichment. Importantly, the wθ score provides an interpretable measure of phenotype robustness, while the statistical significance of each phenotype is captured by the Bayesian-derived, false discovery rate (FDR)-adjusted p-value (Fig. 1C and Suppl Fig. 1).

A further Bayesian statistical test assessed differences in binomial proportion estimates (wθ) between the two conditions (e.g. *in vivo* vs. *in vitro*), using non-targeting control (NTC) clones to set priors (Fig. 1C). To quantify the magnitude of this differential effect between the *in vivo* and *in vitro* conditions, we introduced the weighted delta (wΔ) score, representing the absolute difference in wθ values between the two conditions. This metric captures the extent to which a gene’s phenotype diverges *in vivo* versus *in vitro*. The statistical significance of wΔ scores was evaluated using false discovery rate (FDR)-adjusted p-values providing confidence for this differential score (Fig. 1C and Suppl Fig. 1).

### Validation of CRISPR-StAR for large-scale *in vivo* screening in human tumor xenografts

To optimize the CRISPR-StAR platform for human xenograft models, we first conducted a large-scale pooled CRISPR screen using the non-small cell lung cancer cell line A549 in both *in vitro* culture and in subcutaneous xenograft models (n=118 tumors) (Fig. 1D). The sgRNA library contained 30,000 guides (four guides per gene).

A key feature of CRISPR-StAR is the requirement for balanced recombination outcomes - stochastic activation yielding an active-to-inactive (A/I) ratio near 1 - for effective noise reduction^14^. As such, it is critical that Cre recombination is induced when each clone has expanded enough to generate reproducibility of this stochastic event. *In vivo*, this was achieved by delaying tamoxifen treatment until tumors were established (Suppl Fig. 2A). We confirmed sustained Cas9 expression at the time of recombination by immunohistochemistry (Suppl Fig. 2C) and observed efficient Cre-mediated recombination in the tumors (Suppl Fig. 2D), both prerequisites for robust gene editing.

In the *in vitro* screening arm, we included an intentional bottleneck post-library transduction to mimic the *in vivo* engraftment limitations, followed by a 50-fold expansion prior to 4-OHT induction to empower UMIBB (Fig. 1D and Suppl Fig. 2B). In the *in vivo* arm, aggregating data from all 118 tumors provided comprehensive library representation (Suppl Fig. 3A), with each sgRNA supported by at least 40 distinct clones, which is comparable to the *in vitro* coverage (Suppl Fig. 3B). Interestingly, the number of library-integrated and clonally expanded cells that contributed meaningfully to each tumor plateaued at ∼30,000 clones per sample, irrespective of final tumor size (Suppl Fig. 3C), indicating a potential limit on the number of clones capable of successful engraftment and outgrowth. Injecting 5-fold more or fewer cells did not impact either tumor growth (data not shown) or clonal complexity (Suppl Fig. 3D), further supporting an engraftment bottleneck limitation in this A549 xenograft model.

The CRISPR-StAR screen reliably distinguished core essential genes (ESG; defined as genes essential for viability in > 90% of cell lines in DepMap/Achilles^1^) from non-targeting controls (NTC) and intron-targeting controls (ITC) in both *in vitro* and *in vivo* arms (Fig. 2A). As expected, across multiple experiments, sgRNA dropout was not observed at day 7 post-induction, highlighting the lag time required for recombination and gene editing; thus, we extended the *in vitro* screening endpoints to 21 days and the *in vivo* endpoint to 28 days for the A549 model (Fig. 2A). Despite partial library representation within individual tumors, informative gene dropout data could be extracted using clonal Log_2_A/I ratios. For instance, sgRNAs targeting *PLK1*, an essential gene, consistently showed depletion across tumors, while intron-targeting control sgRNAs (ITC) remained stable (Fig. 2B).

**Figure 2.**
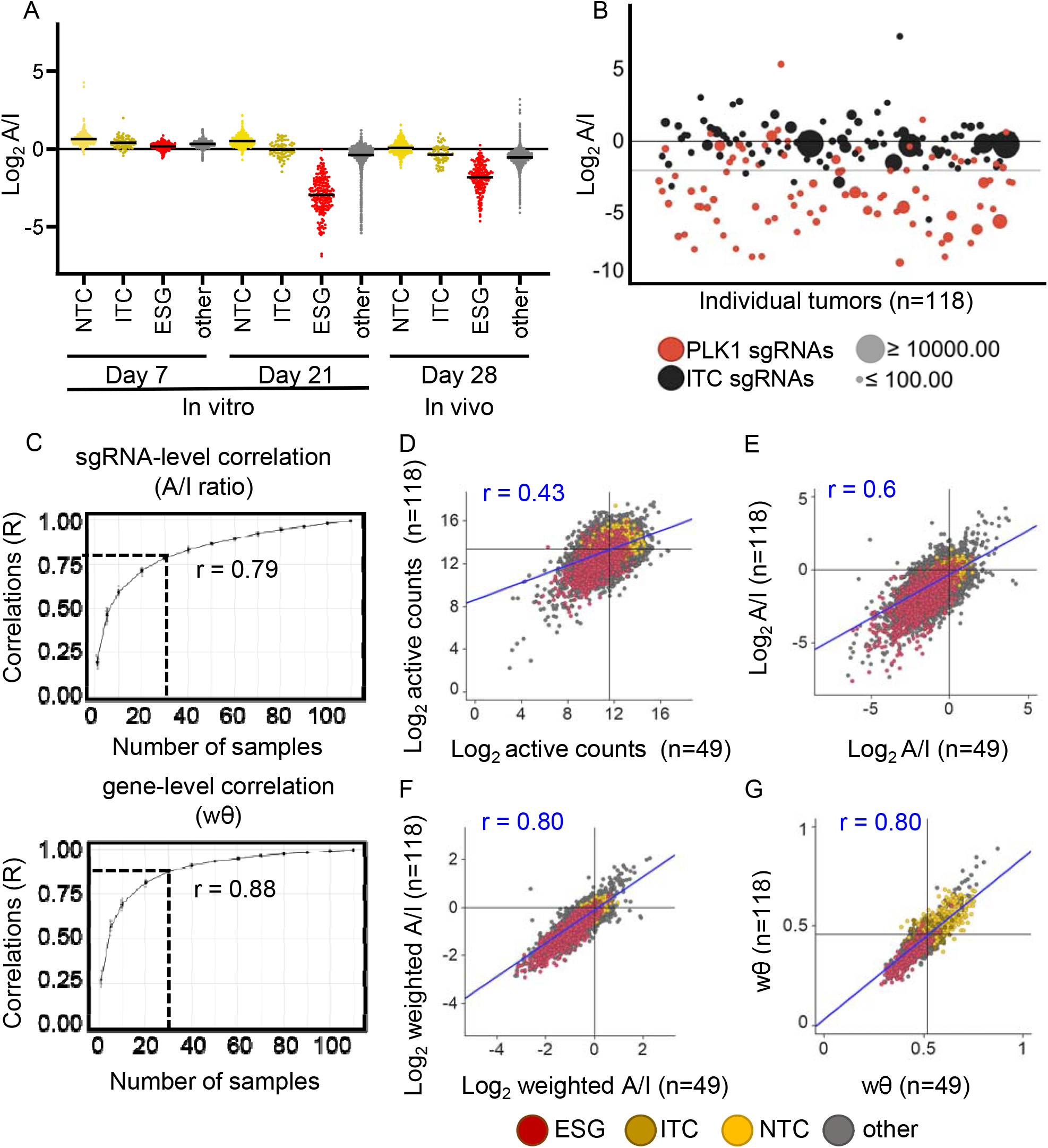
Analysis of 30K sgRNAs CRISPR-StAR screen and downsampling performance. (A) Clonal Log_2_ A/I ratios, averaged across sgRNAs and grouped by gene type, are plotted over time post 4-OHT (*in vitro*) or tamoxifen (*in vivo*) treatment. The library includes 198 essential genes (ESG, 792 sgRNAs), 72 sgRNAs targeting intronic regions (ITC) and 499 non-targeting sgRNAs (NTC), and 6925 other genes (27,700 sgRNAs). (B) Example dropout of the essential gene PLK1 across 118 individual tumors at endpoint compared to intron cutting control (ITC). Each point represents the weighted average log_2_(A/I) ratio per gene per tumor; circle size reflects total read count per gene. (C) Downsampling analysis of *in vivo* data showing sgRNA-level (Log_2_A/I) and gene level (wθ) correlation values between subsets of tumors and the full 118-tumor dataset. X axis: number of tumors in the downsampled subset; Y axis: average correlations of the randomly selected tumor subset versus the full 118-tumor dataset (n=10). Dotted line indicates correlations when 30 random tumors are sampled (r=0.79 for A/I ratio; r=0.88 for wθ). (D-G) Reproducibility of true biological replicates (118 vs 49 mice) for the 30K sgRNA library screen: (D) Correlation of sgRNA-level Log_2_ active counts (r=0.43; proxy for traditional screens without internal controls), (E) Correlation of sgRNA-level Log_2_A/I ratios leveraging internal normalization (r=0.6), (F) Correlation of sgRNA-level Log_2_A/I ratios weighted by clonal size (r=0.8), (G) Correlation of gene-level wθ score from CSTAR-UMIBB analysis (r=0.8).

Although variability in clonal Log_2_A/I ratios was observed – likely reflecting both editing efficiency and NGS limitations – our analysis pipeline effectively distinguished true biological signal from technical noise. By plotting the clonal size (as measured by total sequencing reads per clone) against the Log_2_A/I ratio in each clone, we generated density maps to visualize the overall behaviors of large number of clones (Suppl Fig. 3E). As expected, larger clones exhibited reduced noise, likely due to mitigation of PCR and NGS-related distortions at higher read counts (Suppl Fig. 3E).

### *In silico* down sampling reveals minimal tumor requirement for high-power screening (∼30,000 sgRNAs require 30 tumors)

Given the depth and complexity of our dataset, we hypothesized that our CRISPR-StAR screen may be statistically overpowered and that fewer tumors might suffice to extract reliable sgRNA dropout signals. To test this, we performed *in silico* downsampling of sequencing data from the A549 xenograft screen (n=118 tumors), analyzing both individual sgRNA-level log_2_(A/I) ratios and gene-level wθ scores (Fig. 2C). These analyses confirmed that dropout signals remained highly correlated with the full dataset even with substantial reductions in sample size. Notably, a cohort of just 30 tumors was sufficient to maintain strong concordance with the full 118-tumor screen at both the sgRNA (r=0.79) and gene level (r=0.88), indicating that the CRISPR-StAR system retains high statistical power with drastically reduced animal use compared to conventional xenograft screens that would use hundreds of mice (Fig. 2C).

### Experimental validation of screening depth and reproducibility of CRISPR-StAR

To experimentally validate the above analysis and assess reproducibility across biological replicates, we next performed an independent CRISPR-StAR screen in A549 using just 49 xenograft tumors. We first evaluated the reproducibility of sgRNA dropout based solely on active read counts between biological replicates – a proxy for a conventional CRISPR-UMI^20^ screening without internal controls. This approach yielded low correlation across replicates (r=0.43; Fig. 2D), highlighting the inherent noise in *in vivo* pooled screens^19–23^. Incorporating internal inactive controls for normalization at sgRNA level substantially improved replicate concordance (r=0.60, Fig. 2E). When individual clone size and matched pair normalization were further integrated via the UMIBB framework, reproducibility increased markedly (r=0.80, Fig. 2F), with gene level wθ scores also showing high correlation between replicates (r=0.80, Fig. 2G). Together, these results demonstrate that the CRISPR-StAR platform achieves robust and reproducible sgRNA phenotyping *in vivo*, even with a ∼60% reduction in sample size. Based on clone engraftment rates, down sampling simulations, and biological replication, we estimate that approximately 1,000 sgRNAs can be confidently resolved per tumor in this A549 xenograft model – substantially enhancing screening efficiency while minimizing animal use.

### Identification of *in vivo*-specific phenotypes using CRISPR-StAR

To pinpoint *in vivo*-specific effector genes, we directly compared gene-level wθ scores derived from *in vitro* and *in vivo* screens (Fig. 3A). This analysis identified a distinct subset of genes that promoted tumor growth specifically *in vivo* while having minimal impact on proliferation *in vitro* under standard culture conditions. Applying a stringent significance threshold (ΔFDR <0.001) on the weighted Δ score, we identified 173 genes with differential scores *in vivo* vs *in vitro*. Among these, knockout of 148 genes selectively enhanced tumor growth *in vivo* (Fig. 3B).

**Figure 3.**
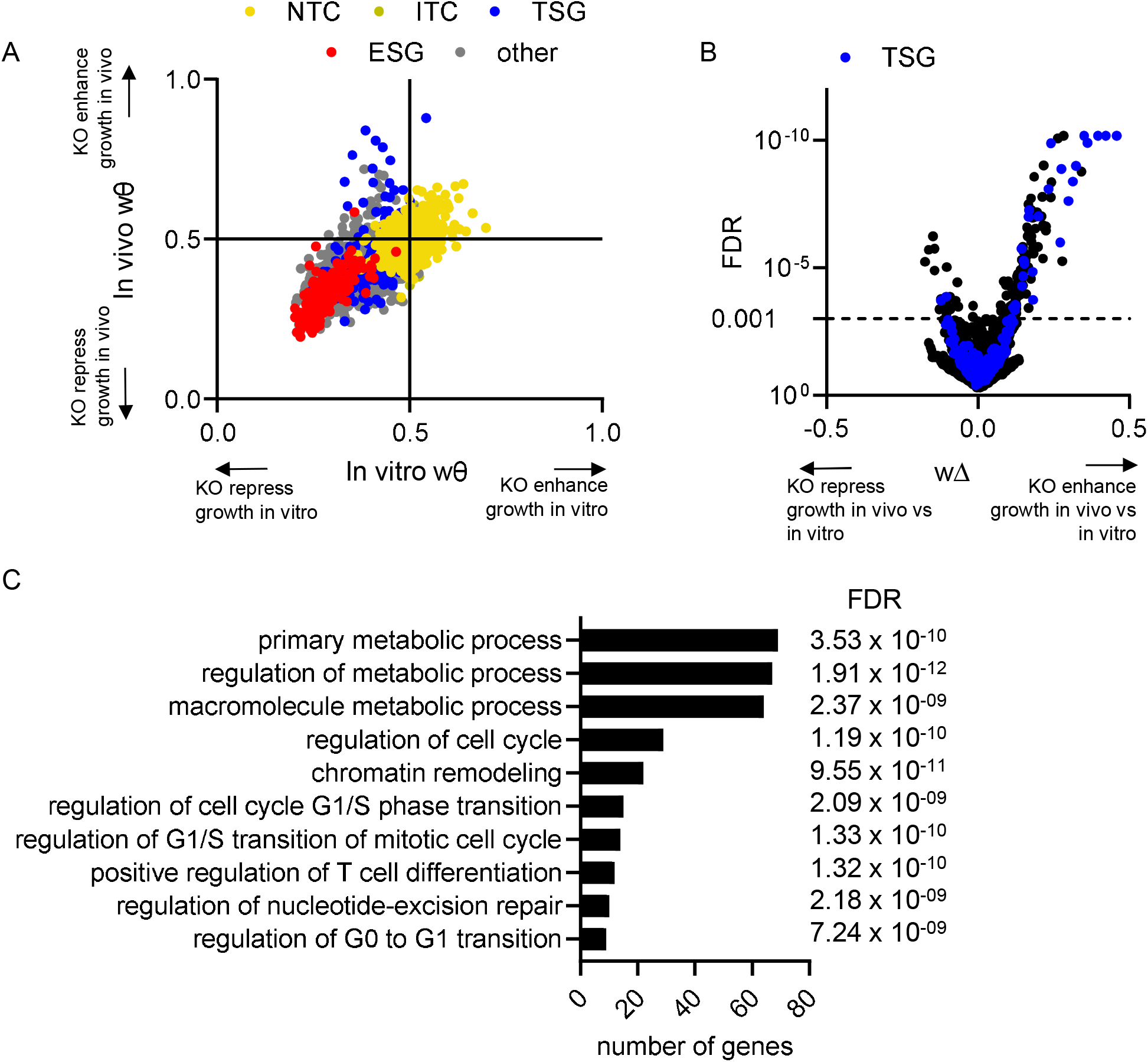
CRISPR-StAR screen identifies *in vivo* specific gene phenotypes. (A) Comparison of gene-level wθ scores from *in vivo* and *in vitro* arms of the 30K sgRNA library screen. Colors indicate sgRNA categories: non-targeting controls (NTC, yellow), intronic controls (ITC, gold), tumor suppressor genes^32^ (TSG, blue), essential genes (ESG, red), and all others (gray). (B) CSTAR-UMIBB analysis comparing *in vivo* vs. *in vitro* phenotypes. X axis: D value attributed by the CSTAR-UMIBB analysis for *in vivo*-specific effect. Y axis: False Discovery Rate (FDR). Dotted line marks FDR=0.001 threshold used to identify 173 hits. (C) Gene Ontology (GO) pathway enrichment analysis of top *in vivo* selective hits from (from panel B), showing selected pathways and associated FDR values. Analysis performed using (https://geneontology.org/)^34,35^.

Notably, this *in vivo* selective gene set was enriched for tumor suppressor genes^32^ with ∼20% of the *in vivo* enrichers categorized as TSGs while only 2.7% of non-significant genes were TSGs (Fig. 3B), which is in line with previous data^33^. Furthermore, there was a marked overrepresentation of epigenetic regulators among the *in vivo* enricher genes (Table 1), including members of the COMPASS family and SWI/SNF chromatin remodeling complexes.

**Table 1:**
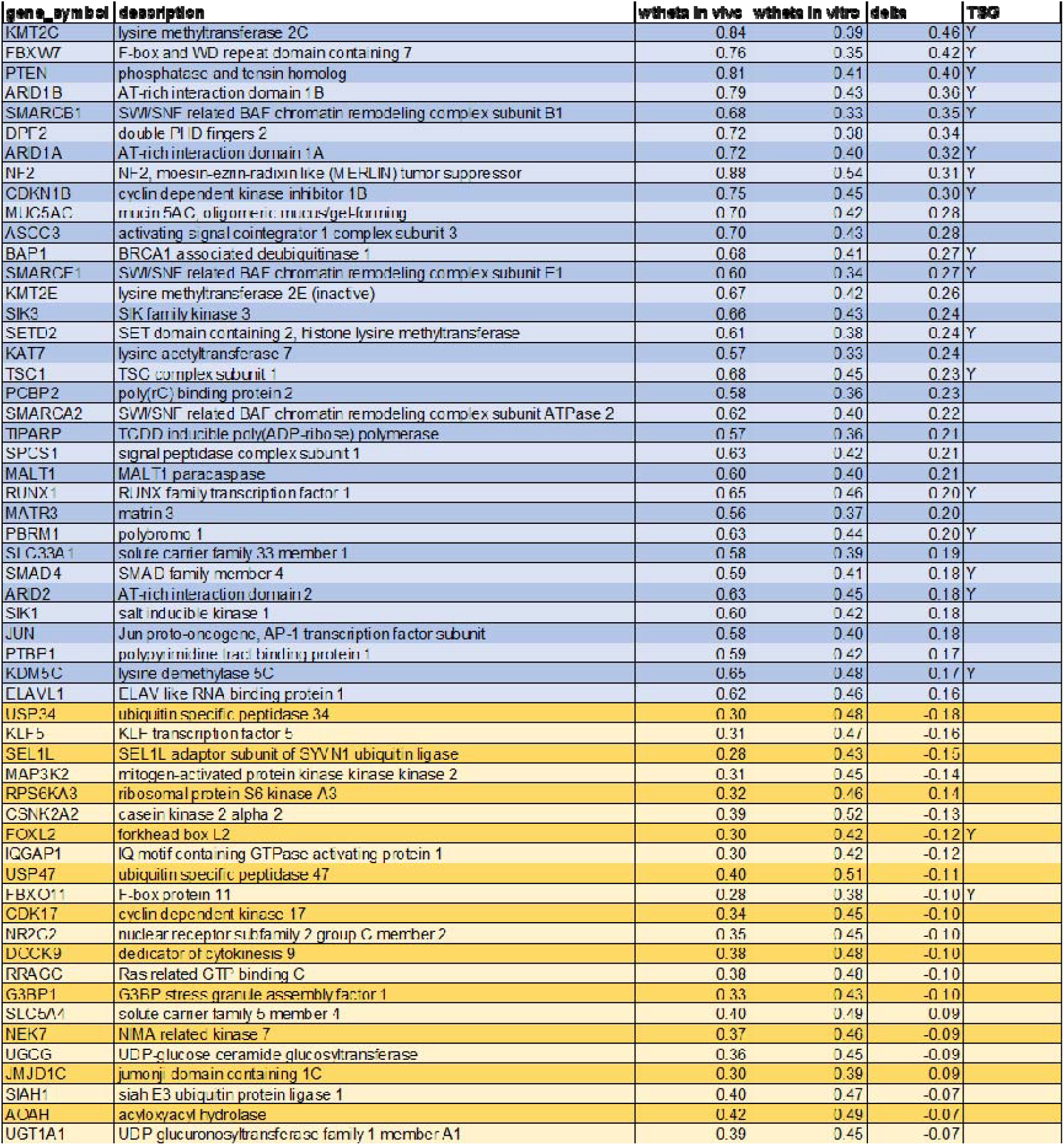
Genes with in vivo selective phenotype upon knockout (in blue are the enriching genes, in yellow the depleting gene).

Gene ontology enrichment analysis^34,35^ confirmed that chromatin organization and remodeling pathways were significantly overrepresented among the *in vivo* specific hits (Fig. 3C), suggesting a key role for epigenetic mechanisms in tumor suppression within physiological *in vivo* tumor environments.

### Mini-pool gene validation confirms *in vivo* specific hits across multiple cancer cell lines

To validate these *in vivo* specific hits across multiple tumor models, and to confirm that a single tumor can functionalize ∼1,000 sgRNAs, we designed a focused validation mini-pool library containing 910 sgRNAs targeting top-ranked genes. This library was cloned into both a traditional constitutive CRISPR-UMI backbone^20^ and the inducible CRISPR-StAR system^14^ (Fig. 4A), allowing head-to-head comparison of validation strategies. While the UMI platform also enables clonal tracking it relies on constitutive sgRNA expression and does not include the matched-pair internal control design unique to CRISPR-StAR^14,20^. The validation screens were performed across three human cancer cell lines (A549, NCI-H460 and SW1573) using comparable number of animals per arm to ensure adequate representation of the UMI library for subsequent bioinformatics analysis. Indeed, across all three xenograft models, both platforms reliably separated core essential genes from non-targeting controls (Suppl Fig. 4A), confirming technical robustness.

**Figure 4.**
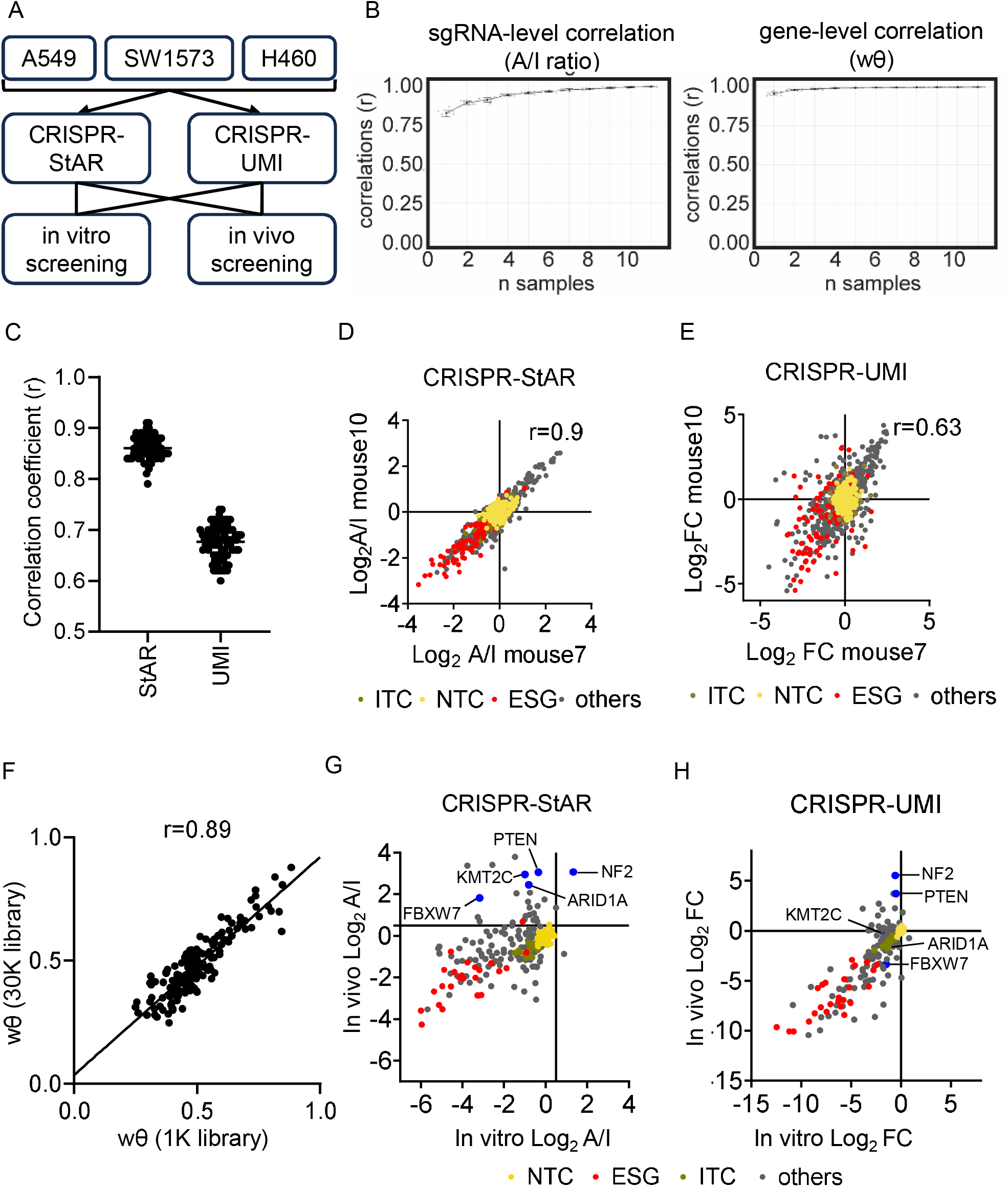
Validation of primary screening hits and comparison of CRISPR-StAR vs. CRISPR-UMI. (A) Schematic of minipool validation screen using a 1K sgRNA library targeting top enrichers and depleters from the 30K screen. Screens were performed in CRISPR-StAR and constitutive CRISPR-UMI systems across three non-small cell lung cancer (NSCLC) cell lines. (B) Downsampling analysis of CRISPR-StAR *in vivo* samples. sgRNA-level log_2_(A/I) and gene-level wθ correlations were computed between subsets and the full 12-tumor dataset. X-axis: number of tumors sampled; Y-axis: average correlation. (C) Average pairwise Pearson correlation (r) of sgRNA profiles across all tumor pairs within each arm (CRISPR-StAR and CRISPR-UMI). Full pairwise plots shown in Supplementary Fig. 3. (D) Representative correlation between two CRISPR-StAR tumors, showing clonal average log_2_(A/I) ratios per sgRNA.(E) Representative correlation between two CRISPR-UMI tumors, showing log_2_ fold change of sgRNA counts between day 28 (study endpoint) and day 0 (injection). (F) Concordance (scatter plot) of gene-level wθ values between the original 30K screen and the 1K validation minipool. Pearson correlation shown. (G) Gene-level comparison of *in vivo* vs. *in vitro* wθ values from the CRISPR-StAR 1K minipool validation screen. (H) Gene-level comparison of *in vivo* vs. *in vitro* fold change from the CRISPR-UMI 1K minipool validation screen.

Importantly, CRISPR-StAR achieved consistent results at the predicted resolution of ∼1,000 sgRNAs per tumor, supported by sufficient clonal coverage (Fig. 4B). Computational down sampling analysis of the dataset showed that a single tumor could recapitulate the signal obtained from a 12-tumor cohort with a correlation coefficient of r=0.96 for wθ and r=0.83 for sgRNA-level A/I ratio (Fig. 4B). Pairwise correlation analysis demonstrated strong reproducibility across individual animals in the CRISPR-StAR arm as compared to the CRISPR UMI arm (Suppl Fig. 4B). This is also evident when the coefficient score of all possible intra-tumor comparisons were summarized in the dot plot (Fig. 4C). As a representative example, the correlation between mouse 10 vs mouse 7 in the CRISPR-StAR arm shows a correlation coefficient of r=0.9 (Fig. 4D), while the correlation of 2 representative animals in the UMI cohort has lower correlation coefficient of r=0.63 (Fig. 4E). These data demonstrate the ability of single-tumor CRISPR-StAR screens to resolve meaningful phenotypes at this scale.

In contrast, although UMI-based data also showed high correlation when the endpoint counts were normalized to plasmid counts (data not shown), closer examination revealed significant pre-transplant drift in sgRNA representation – biases introduced during *in vitro* expansion of the cell pool prior to xenografting due to the *in vitro* gene KO effects. This drift reduced the reproducibility in single tumor UMI screens unless the data were normalized to the pre-injection pool (Suppl Fig. 4C). CRISPR-StAR, by contrast, allows direct *in vivo* assessment without dependence on pre-transplant sgRNA distributions, thereby reducing batch effects and improving signal fidelity.

Unlike conventional CRISPR screening methods, where increased library size can diminish resolution due to competition among sgRNAs, CRISPR-StAR’s internal control structure ensures that sgRNA dropout measurements are independent of library composition. As a result, gene-level dropout signals in the minipool validation screen closely matched those observed in the original 30K-library screen (Fig. 4F), despite differences in sgRNA sequences and library context. No additional enrichment or dropout was observed in the minipool screen, indicating screen saturation. Collectively these findings demonstrate that CRISPR-StAR enables efficient discovery, and reproducible validation of *in vivo*-specific hits across multiple xenograft models and supports high-fidelity functional screening of up to 1,000 sgRNAs per tumor.

### Comparison of *in vivo* specific hits between UMI and CRISPR-StAR libraries

We next analyzed *in vivo*-specific hits identified by both the CRISPR-StAR and UMI-based libraries in A549 xenograft models (Fig. 4G and 4H). Several gene dependencies that were robustly detected using CRISPR-StAR were not captured in the constitutive UMI knockout screen (i.e. *FBXW7, KMT2C, ARID1A*). We attribute this discrepancy to key differences in experimental screening design: specifically, the UMI platform initiates gene KO before engraftment, potentially eliminating cells with early fitness defects *in vitro* that impair their ability to form tumors. In contrast, CRISPR-StAR activates gene knockout after tumors are established, allowing interrogation of functional selection exclusively in the *in vivo* tumor environment.

Furthermore, when this validation screen was extended to three non-small cell lung cancer cell lines, each displayed a distinct set of *in vivo* selective dependencies (Suppl Fig. 5A-B), underscoring the cell-line-specific nature of tumor vulnerabilities and the necessity of context-aware screening strategies. Overall, this comparison highlights the superior sensitivity of CRISPR-StAR for identifying *vivo* gene dependencies that may go undetected with conventional UMI-based *in vivo* or *in vitro* screens.

### Divergent roles of KMT2C and KMT2D in tumor growth revealed by *in vivo* CRISPR-StAR and validated by single-gene knockout studies

To complement our pooled CRISPR-StAR screens, we conducted single-gene target validation focusing on the two chromatin regulators: KMT2C and KMT2D. Both genes encode lysine-specific methyltransferases that function within COMPASS complexes, regulating gene expression programs critical for differentiation and development^36^. Although they are often considered tumor suppressors and functionally redundant^37–42^, our *in vivo* screen in A549 xenografts revealed strikingly divergent phenotypes.

Specifically, KMT2C knockout appeared to consistently enhance tumor growth, indicating a tumor-suppressive function, while KMT2D knockout impaired tumor progression, suggesting it is required for *in vivo* tumor maintenance (Fig. 5A left panel). These phenotypes were strongly supported by opposing wθ values at the gene level (Fig. 5A middle panel) and highly reproducible Log_2_A/I ratio across all four sgRNAs targeting each gene (Fig. 5A right panel). Clonal-resolution analysis of Log_2_A/I ratios further confirmed the trend: KMT2C-KO clones were consistently enriched, while KMT2D-KO clones were depleted across tumors (Fig. 5B).

**Figure 5.**
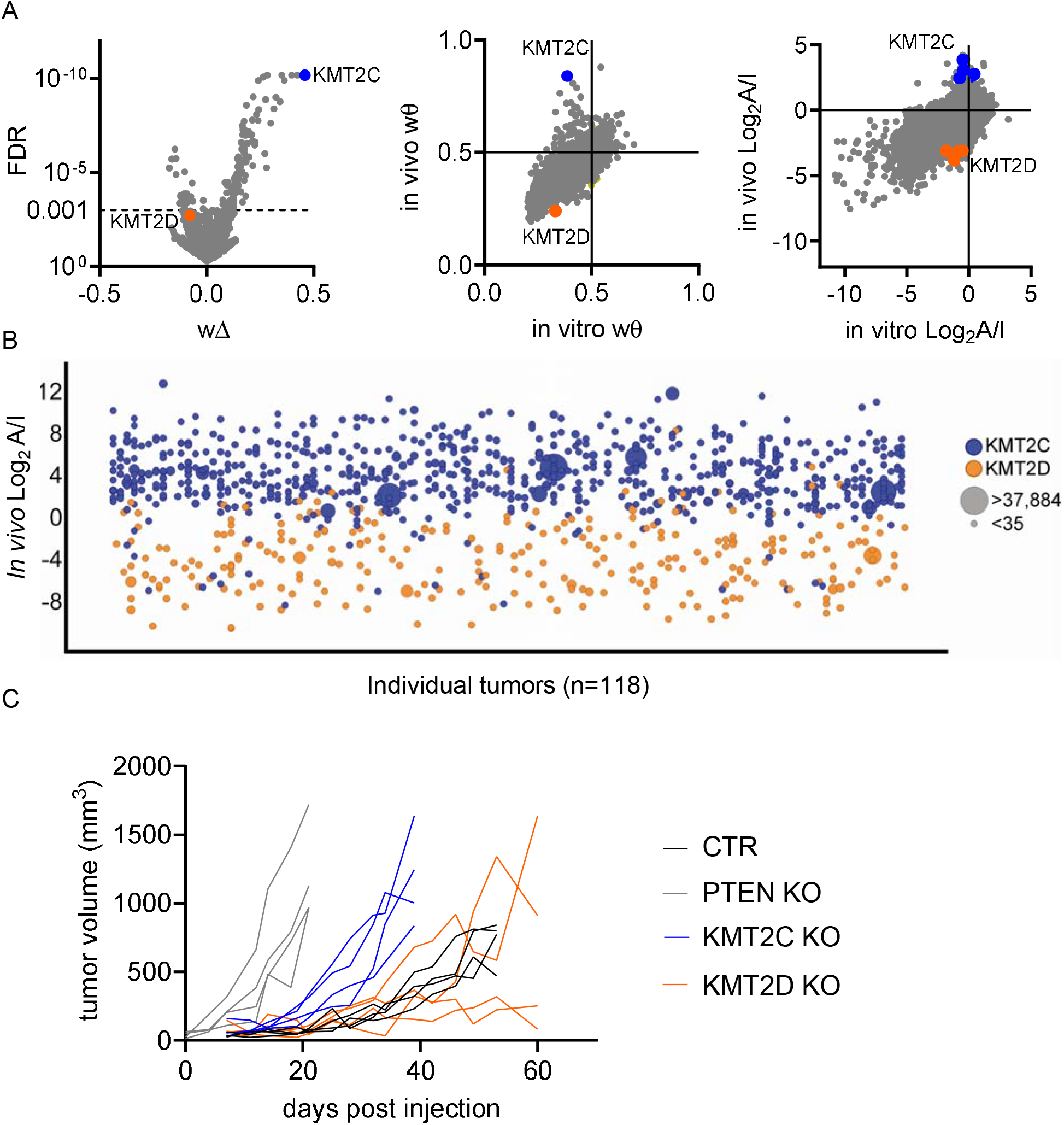
KMT2C and KMT2D have opposite *in vivo* phenotypes in A549. (A) CSTAR-UMIBB analysis highlighting divergent effects of *KMT2C* and *KMT2D*: Left: Δ values (*in vivo* vs. *in vitro*) vs. FDR, with *KMT2C* and *KMT2D* marked. Middle: *In vivo* vs. *in vitro* wθ gene scores. Right: sgRNA-level average log_2_(A/I) ratios *in vivo* vs. *in vitro*, with *KMT2C*/*KMT2D* sgRNAs highlighted. (B) Distribution of log_2_(A/I) ratios for individual clones pertaining to either *KMT2C* or *KMT2D* across tumors. Each point represents a clone; each X-axis position is a separate tumor. Circle size reflects clone size (total sgRNA read counts). (C) *In vivo* tumor growth curves of A549 xenografts with knockout of *KMT2C, KMT2D, PTEN* (positive control), or intron-targeting control (CTR). Tumor volumes plotted individually over time in NCG mice.

To experimentally validate these findings, we generated isogenic A549 cell lines harboring CRISPR-mediated knockouts of KMT2C or KMT2D, or a neutral control sgRNA targeting an intron, as well as a sgRNA targeting PTEN as positive control (Suppl Fig. 6A-D). When these lines were implanted subcutaneously into immunodeficient mice, KMT2C KO accelerated tumor growth, whereas KMT2D KO did not exhibit the same phenotype (Fig. 5C).

Finally, we performed single-cell RNA sequencing (RNA-seq) on time-matched endpoint tumors to validate that underlying transcriptional mechanisms were driving these divergent phenotypes. As expected from COMPASS-mediated enhancer regulation^36^, KMT2C loss led to broad transcriptional changes as compared to intron targeting negative controls with the most differentially expressed genes shown in the heatmap (Fig. 6A). Interestingly, several of the same genes were regulated in the opposite fashion in KMT2D KO tumors (Fig. 6A). This reciprocal expression pattern points to opposing regulatory roles for these methyltransferases in tumor growth and transcriptional control.

Unbiased UMAP (Uniform Manifold Approximation and Projection) clustering of single-cell transcriptomes also revealed distinct tumor cell populations associated with each genotype (Fig 6B). Quantification of the percentage of cells in each cluster as compared to the total pool revealed multiple clusters with distinct representation between KMT2C and KMT2D KO tumors (Fig. 6C). Notably, a cluster of cells (Cluster 7), characterized by high expression of cell-cell interaction genes in control and KMT2D KO tumors, was completely absent in KMT2C KO tumors (Fig, 6B-D). These single-gene validation experiments and transcriptional profiling highlight the power of CRISPR-StAR to detect functional divergent roles *in vivo* for structurally related genes.

**Figure 6.**
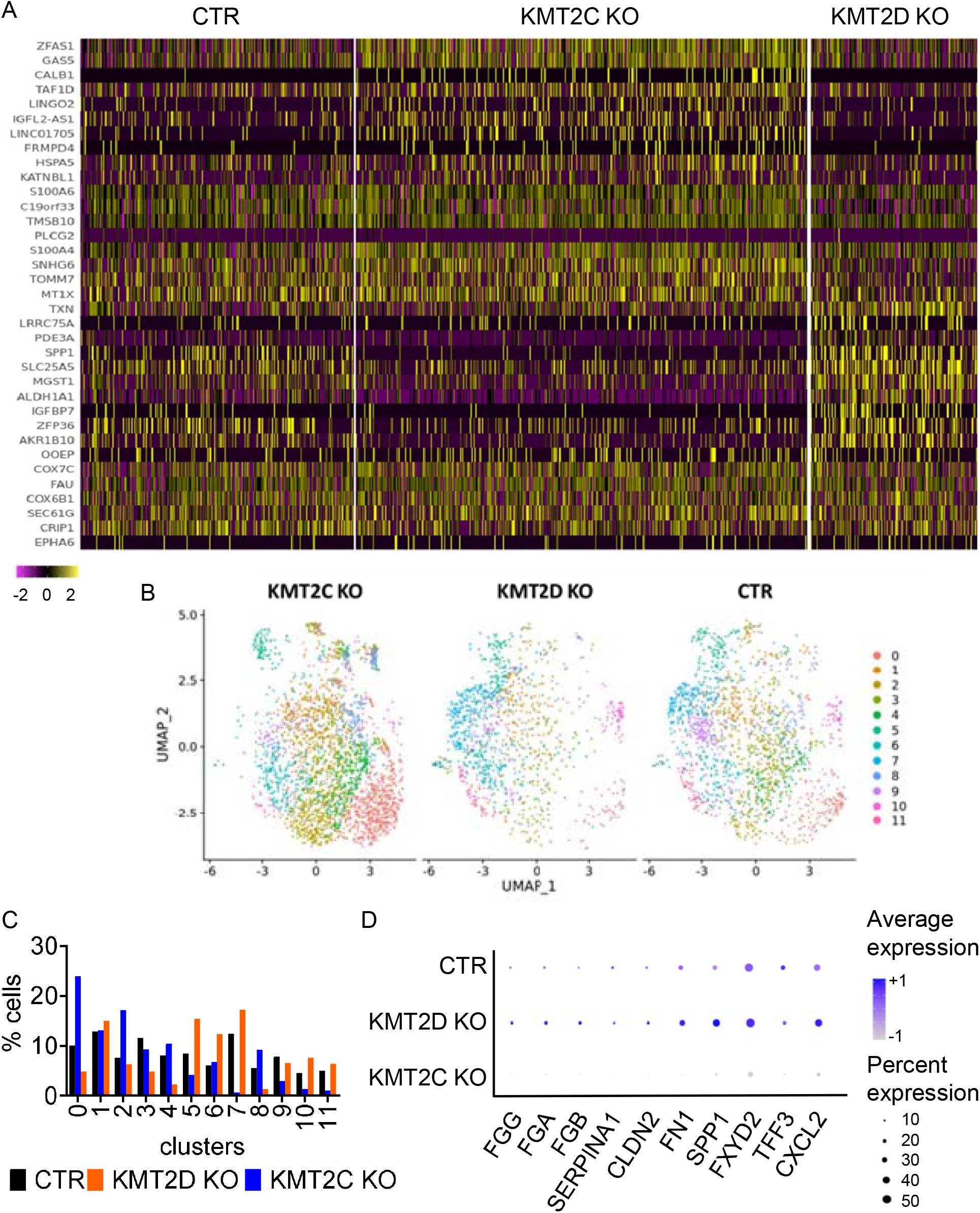
Single-cell RNAseq analysis of KMT2C and KMT2D KO tumors. (A) Heatmap of top differentially expressed genes in KMT2C KO tumor vs. control sample based on single cell RNAseq data (Each block represents one single cell; expression of the same genes is shown for the KMT2D KO tumors). (B) UMAP clustering of single-cell transcriptomes from KMT2C KO, KMT2D KO and control tumors. (C) Quantification of each cluster in shown in panel B expressed as the percentage of total cells per group. (D) Key genes comprising cluster 7 in panel B and C. The average expression of each gene in either control, KMT2D KO or KMT2C KO samples were plotted. Dot size represents the percentage of cells expressing each gene, color indicates average expression level across all cells within each group.

## DISCUSSION

CRISPR-based functional genomics has revolutionized cancer research, enabling systematic identification of genetic dependencies that underlie tumor proliferation, survival and immune evasion. Landmark efforts such as the DepMap and Achilles^1^ projects have mapped essential genes across hundreds of cell lines revealing tumor specific vulnerabilities and synthetic lethal interactions that continue to guide therapeutic development. However, these *in vitro* systems are inherently limited in their ability to recapitulate the complexity of the tumor microenvironment, including cellular heterogeneity, stromal interactions, and nutrient or oxygen gradients - factors that profoundly shape tumor behavior and therapeutic response^13,15^.

Our study addresses this gap by implementing and extending the capabilities of CRISPR-StAR (Stochastic Activation by Recombination) to mouse models of human cancer enabling the activation of gene editing in established xenograft tumors^14^. By temporally separating engraftment from gene perturbation, CRISPR-StAR avoids pre-engraftment selection biases and captures gene dependencies that emerge in the context of the *in vivo* tumor niche. This strategy, combined with our computational pipeline, that implements a Bayesian statistical framework to exploit internally matched sgRNA controls and clonal barcode tracking, enables scripted high-resolution assessment of gene dropout and enrichment scores.

Traditional *in vivo* CRISPR dropout screens suffer from low sgRNA representation and signal-to-noise ratios due to bottlenecks inherent to *in vivo* systems. In cell-line-derived xenograft (CDX) models, less than 2-5% of injected cells contribute to the final tumor, compounded by uneven clonal expansion due to heterogeneous growth within the tumor microenvironment^18^. This leads to low sgRNA representation, typically addressed by increasing the number of animals, an approach that is resource-intensive and, in practice, only partially compensates for the underlying variability.

By contrast, the newly developed CRISPR-StAR (Stochastic Activation by Recombination) is a screening platform tailored for high-resolution *in vivo* screening^14^. CRISPR-StAR achieves improved resolution by (i) activating sgRNAs once tumors are established, and (ii) generating matched-pair internal controls for each clone therefore addressing random variations. Our data builds on the initial application and explores the resolution power of CRISPR-StAR by generating comprehensive single tumor data.

Although existing tools like MAGeCK can be adapted to process CRISPR-StAR data by treating active and inactive clone read counts in CRISPR-StAR as sgRNA abundance in conventional pooled screening^14^, it is only possible when the clonal representation and sample sizes are relatively low. In our experience, the large dataset generated by the CRISPR-StAR platform is over the processing limit of the MAGeCK pipeline to reliably generate results. Furthermore, the discrete clonal read counts may not fit the negative binomial model or other model assumptions held by the existing methods designed based on sgRNA abundance distributions. In addition, the conventional methods are subject to false positive findings due to outlier clones^14^. A prefiltering procedure has been recommended, however, it is often arbitrary to determine how many outlier clones per guide to exclude. By simply comparing counts of active vs inactive control in each clone, the UMIBB approach converts the discrete clonal read counts into depletion/enrichment events count, which can be modeled by a Bayesian binomial distribution. UMIBB approach is also robust to reduce the impact of outlier clones, since the extreme high count in an outlier clone will be converted to just one depletion/enrichment event and won’t have much impact on the gene statistics aggregated on tens or hundreds of depletion/enrichment events. Therefore, the UMIBB approach does not require pre-filtering of outlier clones. Our computational pipeline is an expanded integration of clonal information within the CRISPR-StAR analytical pipeline, improving the robustness of the data by acknowledging the outlier effect.

Our study protocol provided proof-of-concept for the scalability and resolution of CRISPR-StAR. Statistical downsampling analysis followed by validation screens estimated that each individual tumor can resolve ∼1,000 sgRNAs, setting a new efficiency benchmark for *in vivo* genetic screening. Compared to conventional *in vivo* dropout screens, which typically require >100 mice to achieve sufficient library coverage for a 30K sgRNA library, CRISPR-StAR reduces animal use by up to 7-fold without sacrificing statistical power. As such, it makes it a more sustainable and scalable approach for large-scale *in vivo* screens. In our study, we were able to cover approximately one-third of the human genome (30K sgRNA library) with a representation of 40 independent clones (i.e. ‘experiments’) per sgRNAs.

This level of resolution can be achieved because of one of the most advantageous aspects of CRISPR-StAR: its use of inactive sgRNAs as an internal reference, which allows for reliable data extraction even from under-represented screens by focusing on sgRNAs retained in the inactive pool at the endpoint. Traditional CRISPR screening methods lack this reference population and rely on increasing animal numbers to achieve library representation. Additionally, CRISPR-StAR’s internal controls also enhance cross-experimental comparisons. Based on our clone representation and dropout correlation analyses, the reproducibility of true biological replicates yielded highly correlated data across experiments.

A central biological finding of this study is that a substantial subset of tumor suppressor genes exert profound phenotypic effects *in vivo* but remain functionally silent under standard in vitro culture conditions. These genes, frequently mutated or deleted in human cancers, do not influence proliferation in two-dimensional cell cultures, yet significantly accelerate tumor growth upon knockout in xenograft models as exemplified by the KO of PTEN, FBXW7, KMT2C and ARID1A (Fig. 4G and 5C). This highlights a critical and underexplored class of context-dependent tumor suppressors, whose inactivation confers selective advantage only within the physiological constraints of an *in vivo* tumor. Indeed, prior studies have exemplified that certain tumor suppressor genes only manifest a phenotype under in soft agar cultures or *in vivo* conditions^33^. For instance, STK11 is altered in approximately 14% of lung adnocarcinoma^43–46^, however, its genetic knockout (KO) or re-expression does not significantly impact cell proliferation *in vitro* 2D culturing conditions, despite its well-established roles in promoting tumorigenesis *in vivo*^47^.

Notably, our method optimization study revealed 173 genes with divergent *in vivo* vs. *in vitro* phenotypes, including 148 genes whose loss selectively enhanced tumor growth *in vivo*. Many of these genes are involved in epigenetic regulation, particularly within the COMPASS family and SWI/SNF chromatin remodeling complexes, underscoring a role of chromatin dynamics in tumor suppression. Notably, KMT2C and KMT2D, which are paralogous members of the COMPASS complex often considered functionally redundant, displayed strikingly opposing effects: KMT2C loss promoted tumor growth, while KMT2D knockout impaired it. These phenotypes were validated using both single-gene knockout models and single-cell transcriptomic profiling, revealing transcriptional programs uniquely governed by each gene within the *in vivo* tumor ecosystem. Such divergent behavior would likely remain obscured in traditional *in vitro* assays. One of the group of genes identified from our single cell RNAseq analysis are the key factors that regulate cell-cell interactions such as the fibrinogen family genes, which were absent upon KMT2C KO (Fig. 6B-D). Interestingly, FGA KO has been shown to enhance cell migration and invasion in both A549 and other models^48,49^.

Looking ahead, we envision CRISPR-StAR as a foundational tool for building a complementary *in vivo* dependency map to augment the *in vitro* datasets generated by DepMap and Achilles. Such a map would offer deeper insight into tumor specific vulnerabilities, informing the development of therapies tailored to exploit *in vivo* dependencies. Furthermore, because CRISPR-StAR incorporates internal normalization and clonal lineage tracking, it may facilitate meta-analysis across independent experiments, enhancing statistical power and reproducibility across labs and models.

In conclusion, CRISPR-StAR overcomes key limitations of traditional *in vivo* screening approaches by integrating temporal control, internal normalization, and clonal resolution. Our findings underscore the biological and translational value of studying gene function within physiologically relevant environments and provide a compelling case for incorporating *in vivo* functional genomics into future cancer target discovery pipelines.

## MATERIALS AND METHODS

### Cloning of sgRNA and cDNA constructs

All sgRNA and cDNA constructs are built with the 3rd generation lentiviral system. CRISPR-StAR vectors were modified based on previously described contructs^14^ to aid cell line engineering and NGS workflows, with the following changes: (1) a puromycin selection cassette was used instead of the original neomycin cassette, (2) an i7 sequencing adapter was added downstream of the UMI barcode for PCR and illumina sequencing, and (3) PacI restriction sites were added flanking the CRISPR-StAR component to enable enzymatic digestion and enrichment of the target sequences during NGS library preparation. Cre-ERT2 constructs were used as previously described^14^. Individual sgRNAs oligonucleotides were synthesized by Integrated DNA Technologies (IDT, Newark, NJ) and cloned into destination vectors using T4 DNA ligase (New England Biolabs, M0202L). All plasmid constructs were verified by Sanger sequencing at Genewiz (Boston, MA).

### Cell culture and cell line engineering

A549, NCI H460 and SW1573 cell lines were purchased from ATCC (Manassas, VA). All cells were cultured in DMEM (Thermo Fisher Scientific, #11965118) supplemented with 10% FBS and maintained at 37°C under 5% CO_2_. Cell line authentication was performed at LabCorp (Burlington, NC) and routinely tested for mycoplasma contamination using Lonza MycoAlert Detection Kit (Lonza, LT07-318).

To produce lentivirus, transfection mix containing the respective vector, a lentiviral packaging mix (Cellecta, CPCP-K2A), Lipofectamine 3000 (Thermo Fisher Scientific, L30000015) were prepared in Opti-MEM (Gibco, 31985-062) to transfect the Lenti-X cells (Takara Bio, 632180). Fresh DMEM + 30% FBS were used to replace the transfection mix after 16 hours. Viral supernatant was collected 48 hours post-transfection, filtered through a 0.45 µm membrane, aliquoted, and stored at -80 °C.

For infection, lentivirus was added to target cells with 8 µg/mL polybrene (Sigma-Aldrich, TR-1003-G). Media was refreshed overnight. Antibiotic selection was initiated 48 hours post-infection according to manufacturer’s recommended doses.

To generate CRISPR-StAR clones, Cas9 expressing cells were engineered with CRE-ERT2 construct^14^ and seeded by limited dilution for single-cell cloning. Established single-cell clones were screened with CRISPR-SWITCH construct as described previously^50^. The clone exhibiting the highest activity upon 4-OHT treatment with minimal leakage was selected for CRISPR-StAR screen.

### Animal studies

NCG mice were purchased from Charles River Laboratory (strain 572). All animal protocols were approved by the Charles River Institutional Animal Care & Use Committee (IACUC). For tumor inoculation, engineered cell lines were resuspended in Matrigel (Corning, 354263) and PBS (Sigma-Aldrich,) 1:1 solution at a final concentration of 50,000 cells/µL. Each animal was injected with 200 µL cell suspension subcutaneously in the upper flank. Tumor growth and animal welfare were monitored twice a week. Tamoxifen (Sigma Aldrich, T5648) was delivered in corn oil (Sigma Aldrich, C8267. Concentration 100 mg/mL) at a dose of 75 mg/kg by intraperitoneal injection (IP), once a day for 2 consecutive days.

### CRISPR screen

The 30K and 1K library oligonucleotide pools were custom synthesized from Twist Biosciences (San Francisco, CA). The oligo pools were PCR-amplified, purified and cloned into destination vectors via Golden Gate Assembly with Esp3I (NEB, R0734L). The ligated library plasmids were purified using spin columns (Zymo Research, D4014) and electroporated into Endura ElectroCompetent cells (Bioresearch Technologies, 60242-2). Following recovery, transformed cells were plated onto 245 mm2 bioassay agar dishes (Teknova, L6010) and incubated overnight incubation at 32 °C. Colonies were scrapped into fresh LB medium and pelleted by centrifugation. The final plasmid library was prepared using Qiagen Maxiprep Kits. The plasmid pool was then used for transfection utilizing 2-layer CellSTACK vessels (Corning, 3313). Viral supernatants were aliquoted and stored in -80 °C freezer until further use.

To perform the 30K CRISPR-StAR screen, the A549 CRISPR-StAR clone was infected at low MOI at library representation of 1000x. In the *in vitro* arm, an initial bottleneck at 50X was applied post-library transduction to simulate clonal selection pressure observed *in vivo*. This was followed by a 50-fold expansion before recombination was induced with 4-hydroxytamoxifen (4-OHT). Cells were cultured for 21 days before to endpoint sample collection. For the *in vivo* arm, the library infected A549 CRISPR-StAR cell pool was expanded without bottlenecking and injected subcutaneously into 118 mice at 10 million cells per mouse. Tamoxifen treatment was initiated once tumors reached 150mm^3^. Tumors were allowed to grow for 28 days before mice were euthanized and tumors harvested.

Genomic DNA was extracted from both *in vitro* and *in vivo* samples and subjected to PCR amplification targeting regions surrounding the CRISPR-StAR components. Amplified PCR products were purified with KAPA beads and sequenced on an Illumina NextSeq 2000 platform.

### CRISPR-StAR screen NGS sequence alignment and read count normalization

FASTQ files are mapped to the reference file consisting of both guide sequences and barcode sequences using custom scripts. After mismatches and low-quality reads are removed, only aligned reads are used to measure the abundance of UMIs with an active or inactive form of construct respectively. Very small clones with less than 10 reads (active and inactive combined counts) are removed from further analysis. A pseudo count of 0.5 is added when the active or inactive count is 0.

### CRISPR-StAR screen normalized A/I ratio defines clonal depletion or enrichment events per guide

In each sample, the active count and inactive count of each clone is normalized by the ratio between the sum of counts of the active non-targeting control (NTC) guides and the sum of counts of the inactive NTC guides. A/I Ratio (*R*_*si*_) of the normalized active count (*n*_*Asi*_) vs inactive count (*n*_*Isi*_) of the same clone UMI(*i*) of sgRNA (*s*) is calculated to determine if the active form guide was enriched (*R*_*si*_ > 1) or depleted(*R*_*si*_ < 1) comparing to the inactive form of the same clone. Guide level log2(A/I ratio) was derived to show the effect sizes by averaging of log_2_(*R*_*si*_) weighted by combined counts of each clone, then normalized by the median log_2_(*R*_*s*_) of guides targeting essential genes.

### Unique Molecular Identifier Bayesian Beta-binomial (UMIBB) analysis for UMI-CRISPR pooled screening data

We applied the UMIBB analysis previously reported by Simoneau A.,et.,al.^51^ in the pooled screening dataset of A549 cell line. Briefly, the counts in case group vs control group of the same UMI clone are compared to determine the number of depletion or enrichment events per sgRNA, Then the probability of depletion or enrichment for each sgRNA is modeled by a beta-binomial distribution and testing in a Bayesian framework.

### UMIBB analysis for one group CRISPR-StAR samples

Let be the total distinct clones of sgRNA (*s*), *E*_*s*_ is the number of enriched clones (*R*_*si*_ > 1). It can be modeled as Bernoulli trials process follows beta-binomial distribution with a conjugate prior,

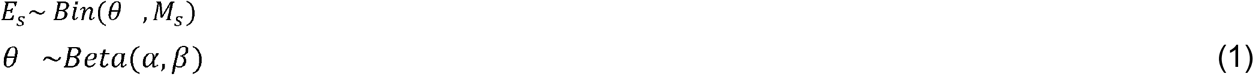

where is *θ* the prior probability of active count greater than inactive count in a clone. The prior probability θ and parameters α, β are estimated using the observed numbers of enriched clones (*E*_*NTC*_) across all the clones with NTC guides (*M*_*NTC*_). The prior θ is equal or very close to 0.5, because the NTC clones are expected to have same chance of enrichment or depletion if there is no systematic bias. Markov chain Monte Carlo (MCMC) approximation is used to fit the posterior probability distribution of *θ*, which combined the prior and the likelihood functions using Bayes’ theorem. The posterior estimate (θ_*Es*_) for clones showing an enrichment of active form measures the effect size for the sgRNA. P-value (*p*_*θEs*_) is estimated by MCMC to test the null hypothesis that *θ*_*Es*_ is less than or equal to the mean prior (θ_*NTC*_). Similarly, we build beta-binomial models using the number (*D*_*s*_) of depleted clones (*R*_*si*_ > 1) and *M*_*s*_, then derived the posterior estimate (*θ*_*Ds*_) for clones showing a depletion of active form and associated p-value (*p*_*θDg*_). Then, the gene level depletion test statistics (*wθ*_*Dg*_,*p*_*θDg*_) and enrichment test statistics (*wθ*_*Dg*_,*p*_*θDg*_)are calculated using the weighted Stouffer’s Z-score method^52^ [ref. 2] weighted by the number of distinct clones in each sgRNA(*M*_*s*_) targeting gene (g). Because most clones are either depleted or enriched for the active form, only very few have equal number of active and inactive form, the sum of *wθ*_*Eg*_ and *wθ*_*Dg*_ is equal or very close to one. So we further consolidate the statistics by using the lower p-value from the depletion and enrichment tests as the gene-level p-value. The relative frequency of clonal enrichment frequency (*wθ*_*g*_) is defined by equation (2), which is plotted as the effect size of genes in the volcano-plot. Finally, the gene-level p-values are adjusted for the multiple testing by the False Discovery Rate (FDR) procedure.

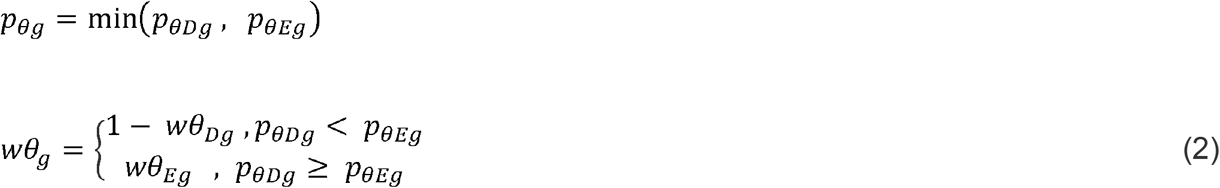

### UMIBB analysis comparing CRISPR-StAR samples between two experiment groups

To compare the difference of active form guide enrichment probability between two experiment groups, we use two independent beta-binomial models to fit the observed number of clones (*E*_*s*1_,*M*_*s*1_) and (*E*_*s*2_,*M*_*s*2_) in the two experimental groups in the same way as defined previously in the single group analysis.

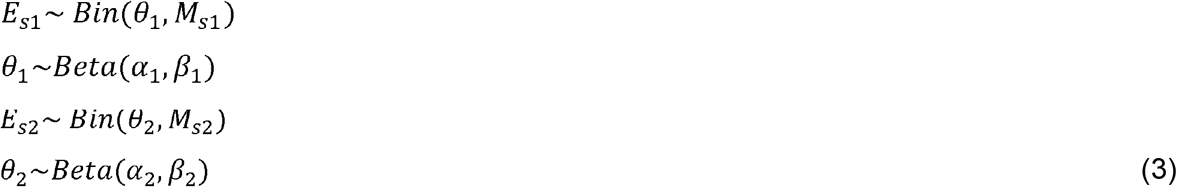

The priors *θ*_1_ and *θ*_2_ are estimated based on the number of active enriched NTC clones (*E*_*NTC*1_,*M*_*NTC*1_) and (*E*_*NTC*2_,*M*_*NTC*2_) among NTC clones in each group separately(3). The posterior estimates (*θ*_*Es*1_,*θ*_*Es*2_) are derived by Bayesian inference as describe before.

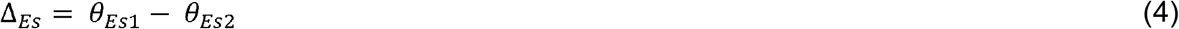

The Δ_*Es*_ is the difference between the posterior proportion estimates of the two experimental groups(4). P-value (*p*_Δ*Es*_) is estimated by MCMC to test the null hypothesis that Δ_*Es*_ is equal to zero.

Similarly, we fit the observed number of clones (*D*_*s*1_, *M*_*s*1_) and (*D*_*s*2_, *M*_*s*2_) for their depletion probability in the two experimental groups, and tested the null hypothesis that the difference (Δ_*Ds*_) of their posterior estimates (*θ*_*Ds*1_, *θ*_*Ds*2_) is equal to zero (5).

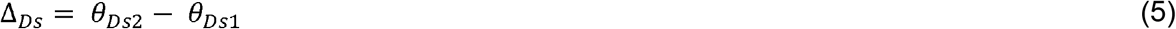

Then, the gene level depletion test statistics of two groups (*w*Δ_*Dg*_,*p*_Δ*Dg*_) and enrichment test statistics (*w*Δ_*Dg*_,*p*_*θDg*_)are calculated using the weighted Stouffer’s Z-score method weighted by the number of distinct clones in each sgRNA targeting gene (g) combined across the two groups. The gene level statistics were consolidated to use the greater significant value of (*p*_Δ*Dg*_, *p*_Δ*Eg*_) and its associated *w*Δ value for the gene (6). *w*Δ_*g*_ is the difference of posterior estimates of clonal enrichment probabilities between the two experimental groups, the p-values (*p*_Δ*g*_) of gene-level tests are adjusted by the FDR procedure.

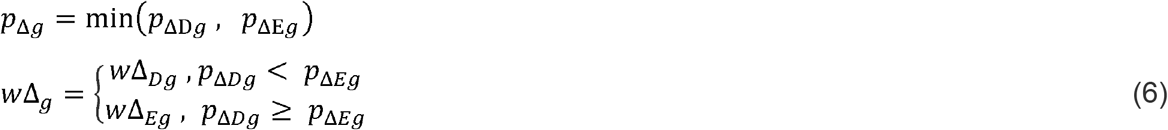

### Animal sequencing counts data down-sampling analysis

Down-sampling of the original full experimental dataset was performed by the sample() function without replacement in R. We randomly took 10 subsets of down-sampling samples for each sample size (n animals) group and calculated the statistics (log2(A/I ratio),*wθ*_*g*_) for each subsets individually. Pearson correlation coefficients (R) are calculated against the statistics of the original full experimental dataset.

### Single cell RNAseq

Tumors were collected from either control A549 or KMT2C or KMT2D KO xenograft models and subjected to 10x genomic protocol for tumor dissociation for single cell RNAs sequencing (protocol#CG000147). Dead cells were removed from single cell suspensions following 10x genomic instruction (protocol#CG000093). Final prepared single cell suspension was then processed with Chromium Next GEM Single Cell 3□ Reagent Kits v3.1 following the manufacturer’s protocol (#CG00031) and subjected to Illumina sequencing on Nexseq 2000.

### Single cell RNAseq analysis

The fastq files from 10x Chromium were demultiplexed and mapped with Cellranger v6.1.2 using a GRCh38 reference provided by 10x Genomics. The feature barcode matrix was then loaded into R for quantification and normalization by the Seurat v4.3 package^53^. Cells classified as negatives or doublets were removed. Cells with more than 2500 genes or fewer than 200 genes were removed, along with cells with a mitochondrial expression fraction above 0.05. Cell cycle effects were calculated using *CellCycleScoring* and regressed out with *ScaleData*. Differentially expressed genes were identified using the MAST package^54^, with the top ones visualized using *DoHeatmap*. We performed graph based Louvain clustering with *FindNeighbors* and *FindClusters* on dimensionality reduced data from *RunPCA* and *RunUMAP*, visualizing 12 Louvain-identified clusters. Percent cell expressions were determined using the *DotPlot* function.

## Availability of data and materials

Additional information can be requested by contacting the corresponding author. The CSTAR-UMIBB source code is packaged into an R-library and available at https://github.com/tangotx/CRISPRstar

## Acknowledgements

This study is funded by Tango Therapeutics.

## FIGURE LEGENDS

**Supplementary Figure 1.**
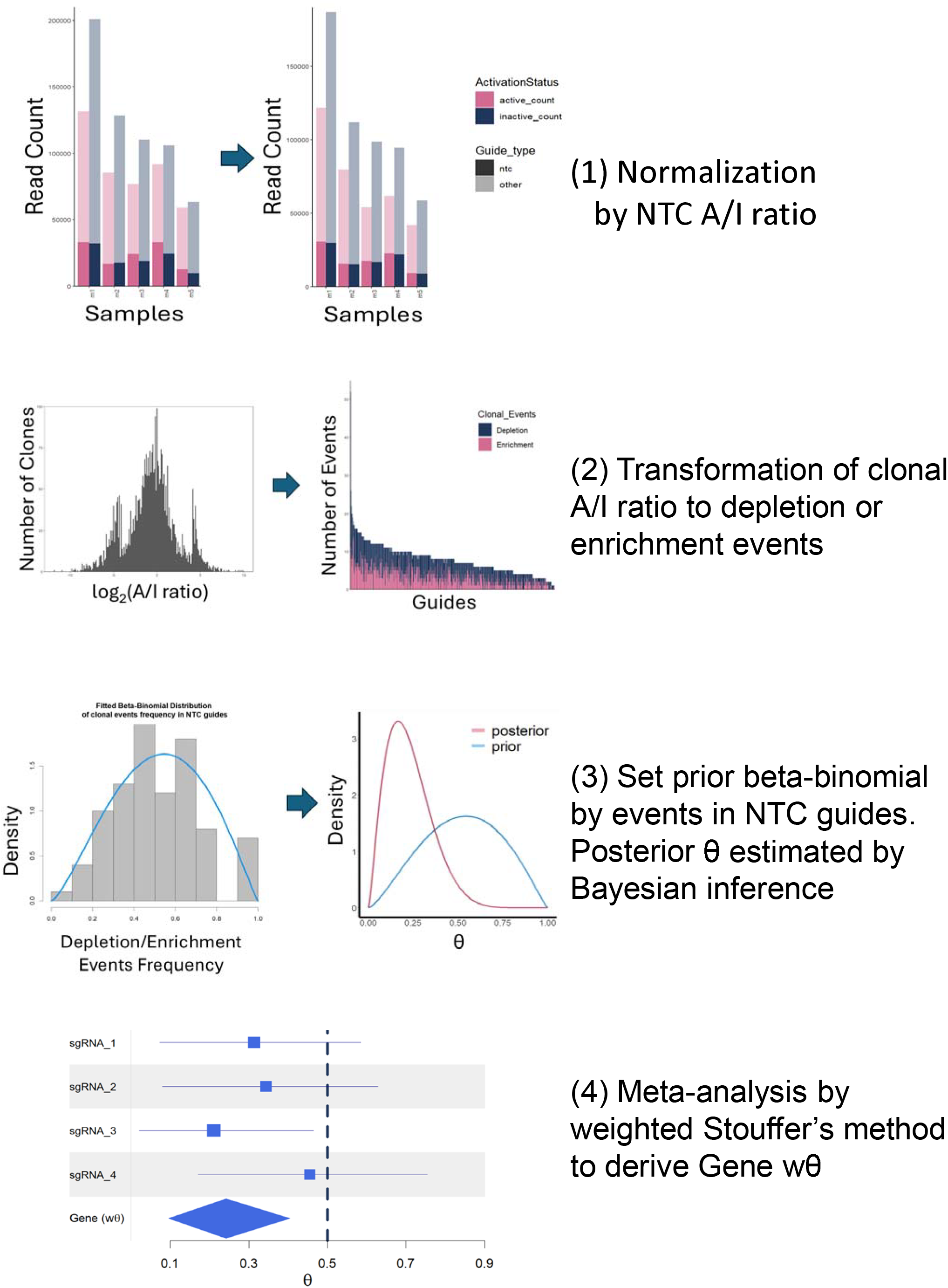
Overview of CSTAR-UMIBB workflow. Schematic depiction of the CSTAR-UMIBB workflow for analyzing CRISPR-StAR screen results.

**Supplementary Figure 2.**
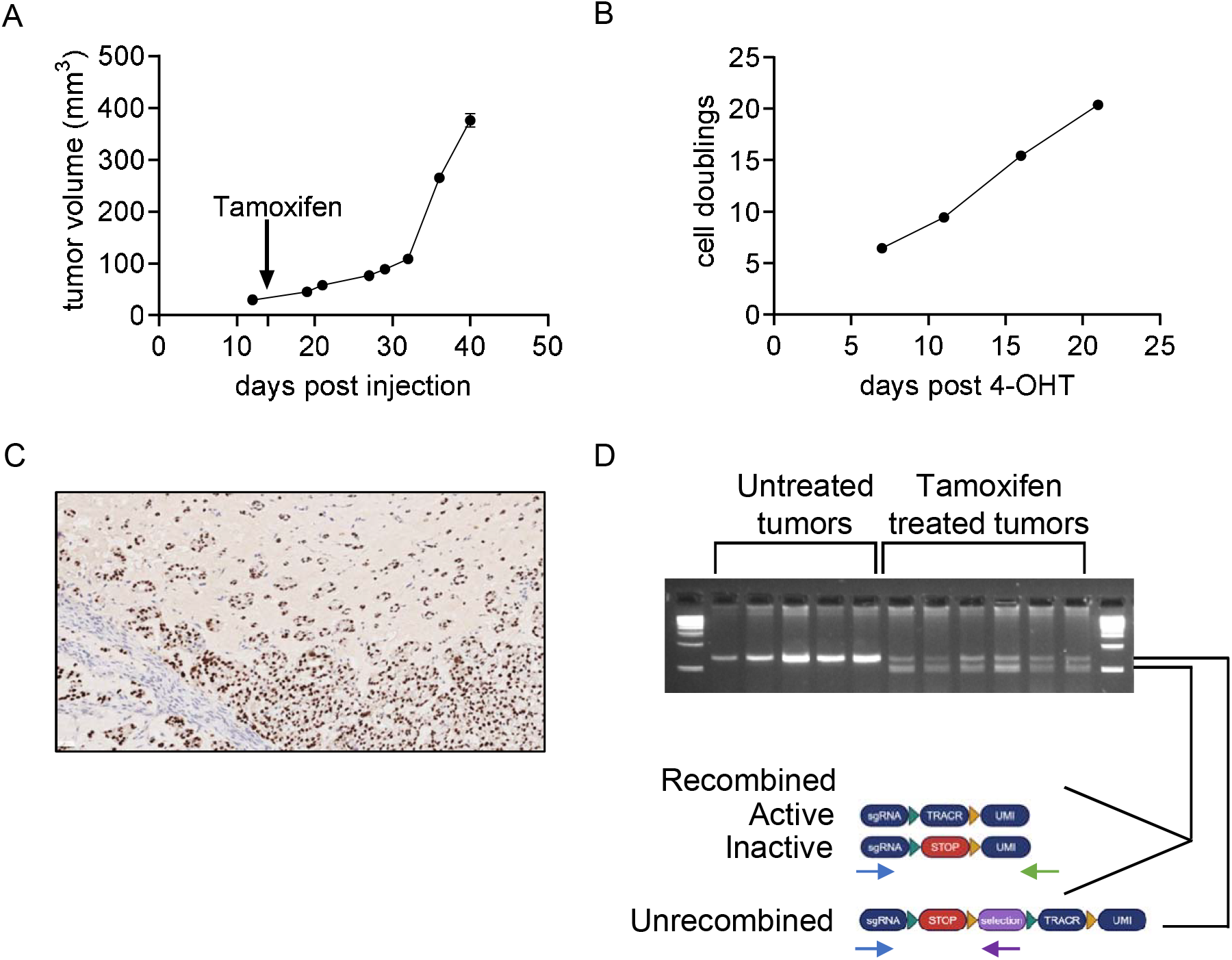
*In vitro* and *in vivo* performance of the 30K sgRNA CRISPR-StAR screen in A549 cells. (A) *In vivo* tumor growth curve (average, n=118) from the 30K sgRNA library screen. Tamoxifen administration was initiated as indicated on day 14 post-injection. (B) *In vitro* cell proliferation measured as population doublings following 4-OHT treatment, starting from day 1 (C) Cas9 immunohistochemistry (IHC) staining in the tumor sections collected 10 days postinjection. (D) Assessment of CRISPR-StAR vector recombination in the tumors 7 days after tamoxifen treatment.

**Supplementary Figure 3.**
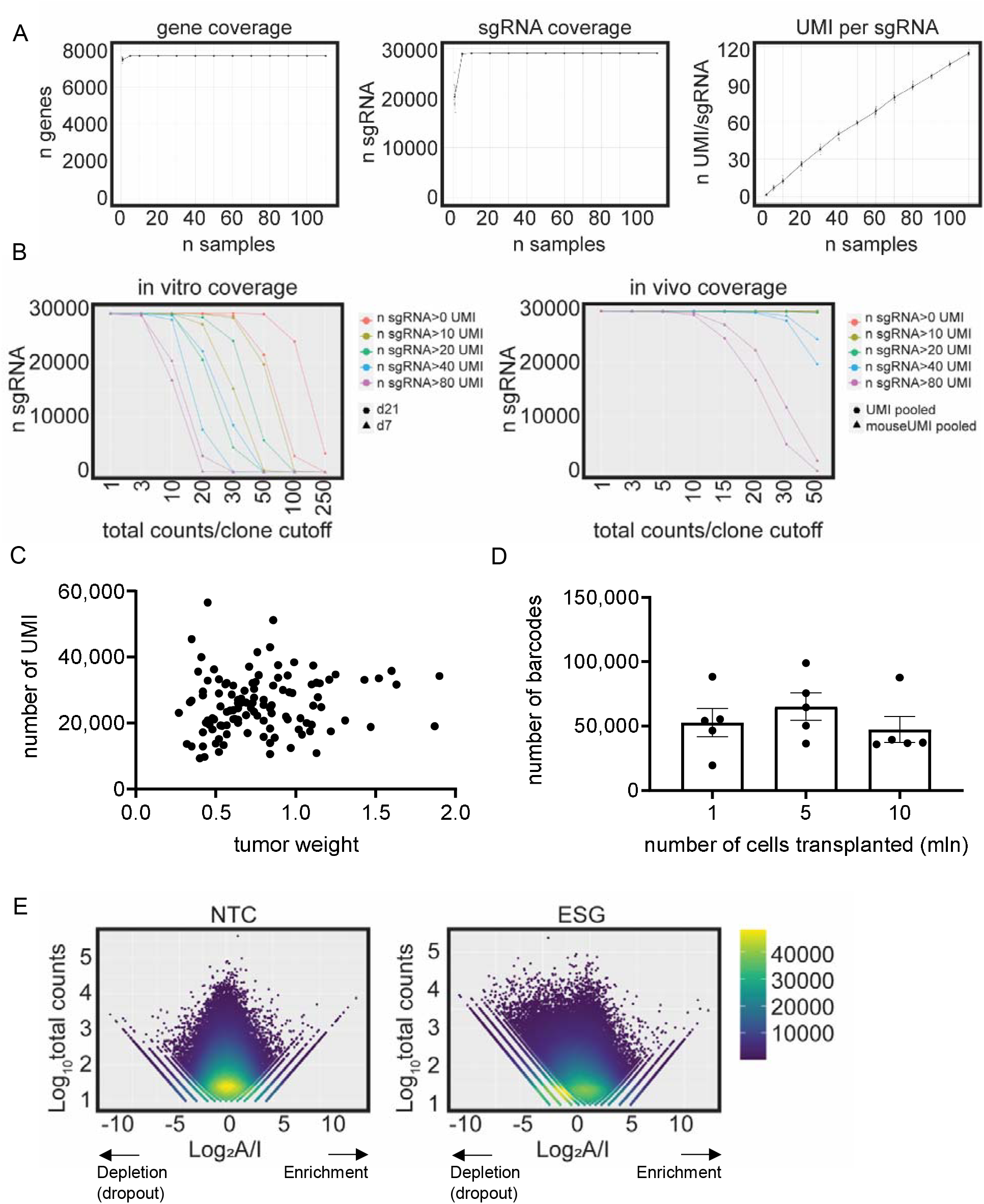
Bioinformatic analysis of of the 30K sgRNA CRISPR-StAR screen in A549. (A) Library coverage metrics across tumor samples: Left panel: gene coverage as a function of tumor number. Middle panel: sgRNA coverage as a function of tumor number. Right panel: Average UMI counts per sgRNA across tumors (B) sgRNA recovery *In vitro* and *in vivo* at varying read count thresholds. X axis: minimum read count per clone. Each clone is identified as an individual sgRNA with unique UMI; Y axis: number of recovered sgRNAs. Colored lines represent different UMI count cutoffs per sgRNA. (C) Correlation between total number of UMI counts vs. tumor weight for each sample. (D) UMI barcode recovery following injection of 1 million, 5 million or 10 million cells. (E) Density plot of clones with either negative control sgRNAs (NTC) or essential gene targeting sgRNAs (ESG). X axis: Log_2_A/I ratio. Y axis: Log_10_ total sequencing counts per clone.

**Supplementary Figure 4.**
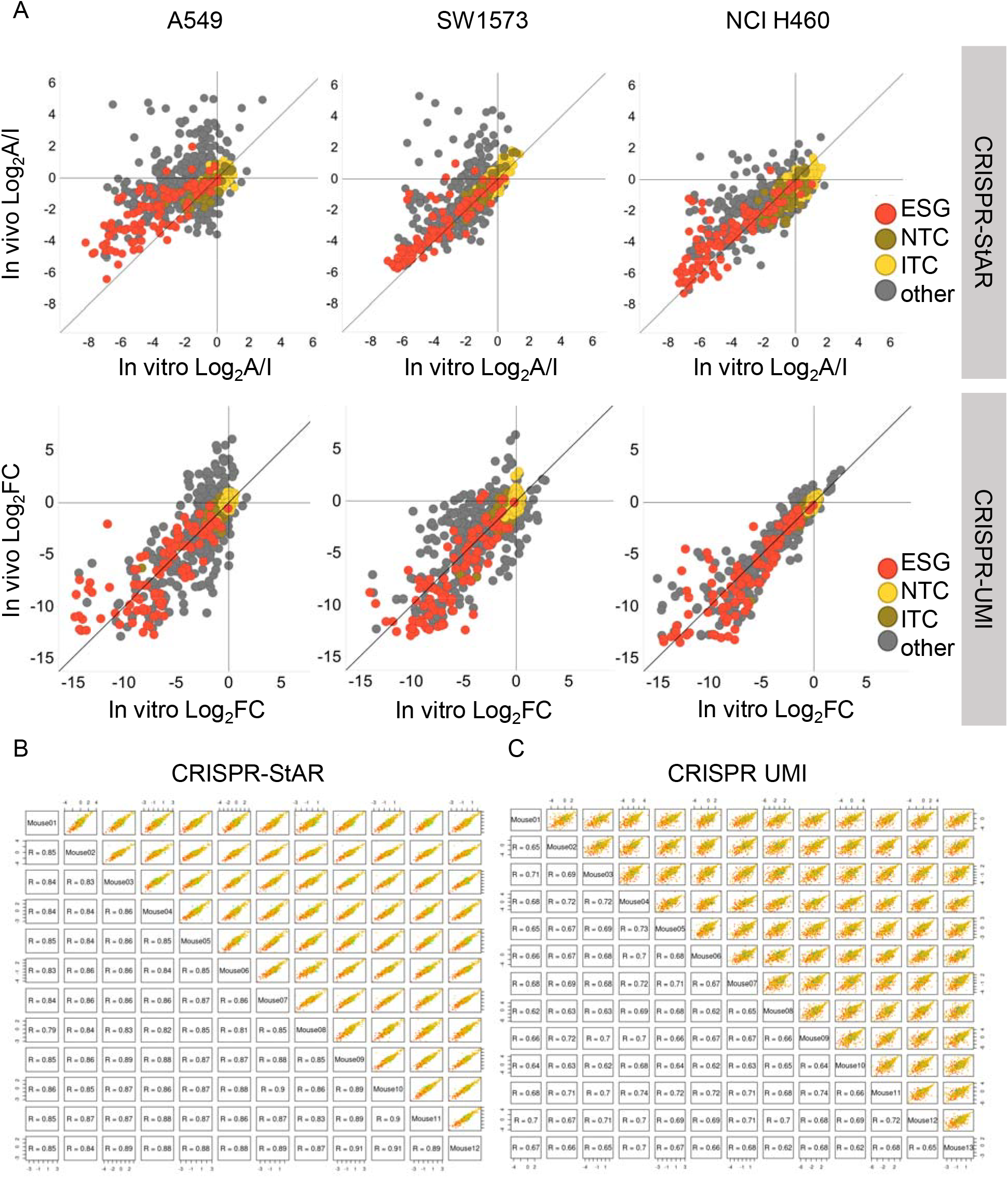
Bioinformatic comparison of constitutive CRISPR-UMI and inducible CRISPR-StAR minipool screens. (A) *In vivo* vs. *In vitro* effect at the sgRNA level of the 1K minipool validation screen across indicated cell lines. For CRISPR-StAR, average Log_2_A/I ratios per sgRNA are shown; for CRISPR-UMI screen, average fold changes of endpoint counts normalized to plasmid counts per sgRNA are shown. (B) Pairwise correlation of sgRNA-level profiles across tumor samples. For CRISPR-StAR screen, average Log_2_A/I ratios per sgRNA were used; for CRISPR-UMI screen, average fold change of endpoint counts normalized to injected pool counts were used for each sgRNA plotted.

**Supplementary Figure 5.**
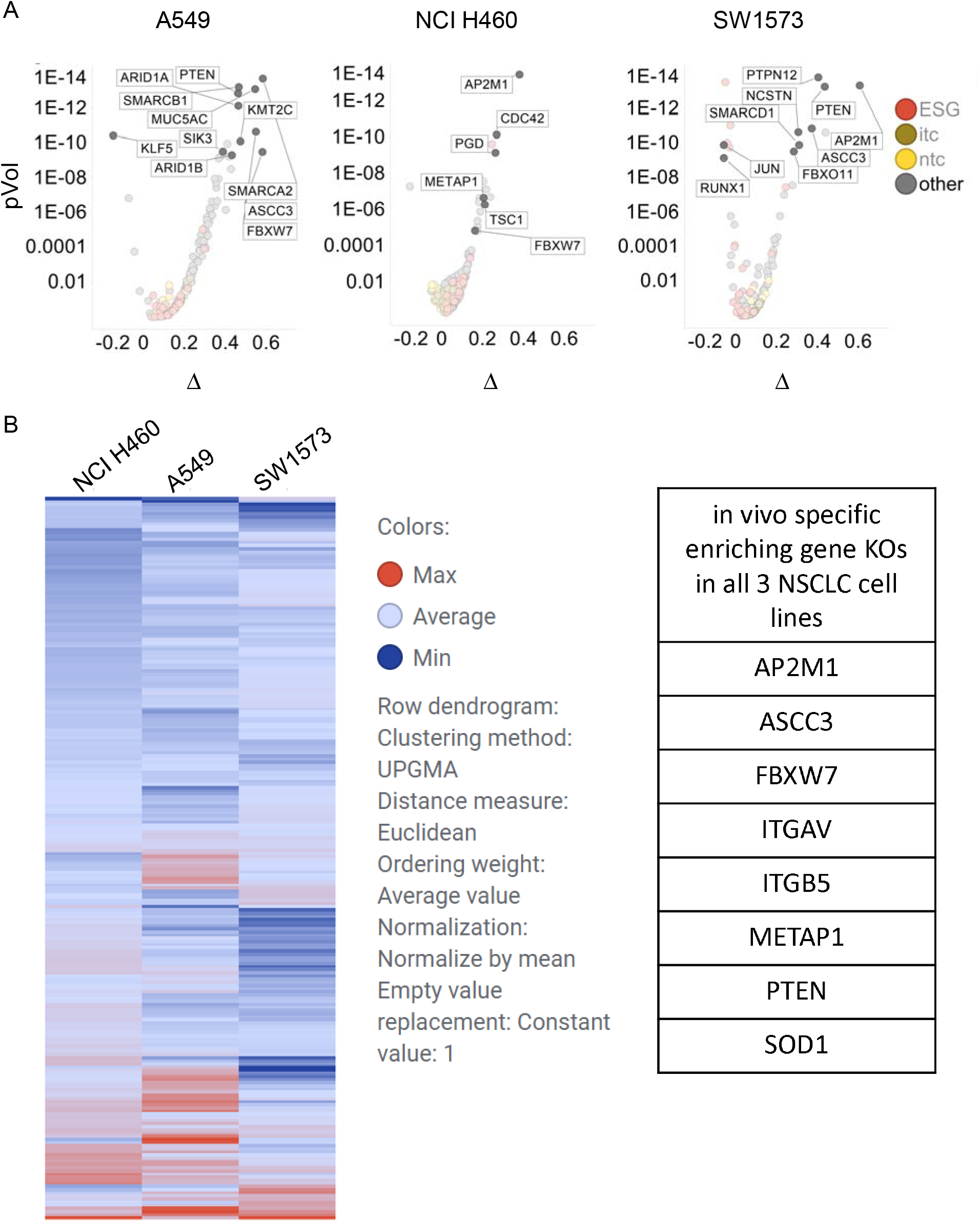
CSTAR-UMIBB analysis of minipool validation library screen across three 3 CDX models. (A) CSTAR-UMIBB gene-level analysis comparing *in vivo* vs. *in vitro* effects for the minipool validation screen in each CDX model. Top genes showing in vivo enriching phenotype are highlighted. (B) Heatmap of top *in vivo* enriched hits per CDX model. Gene shared across all three models are listed in the accompanying table Supplemtary Table 1.

**Supplementary Figure 6.**
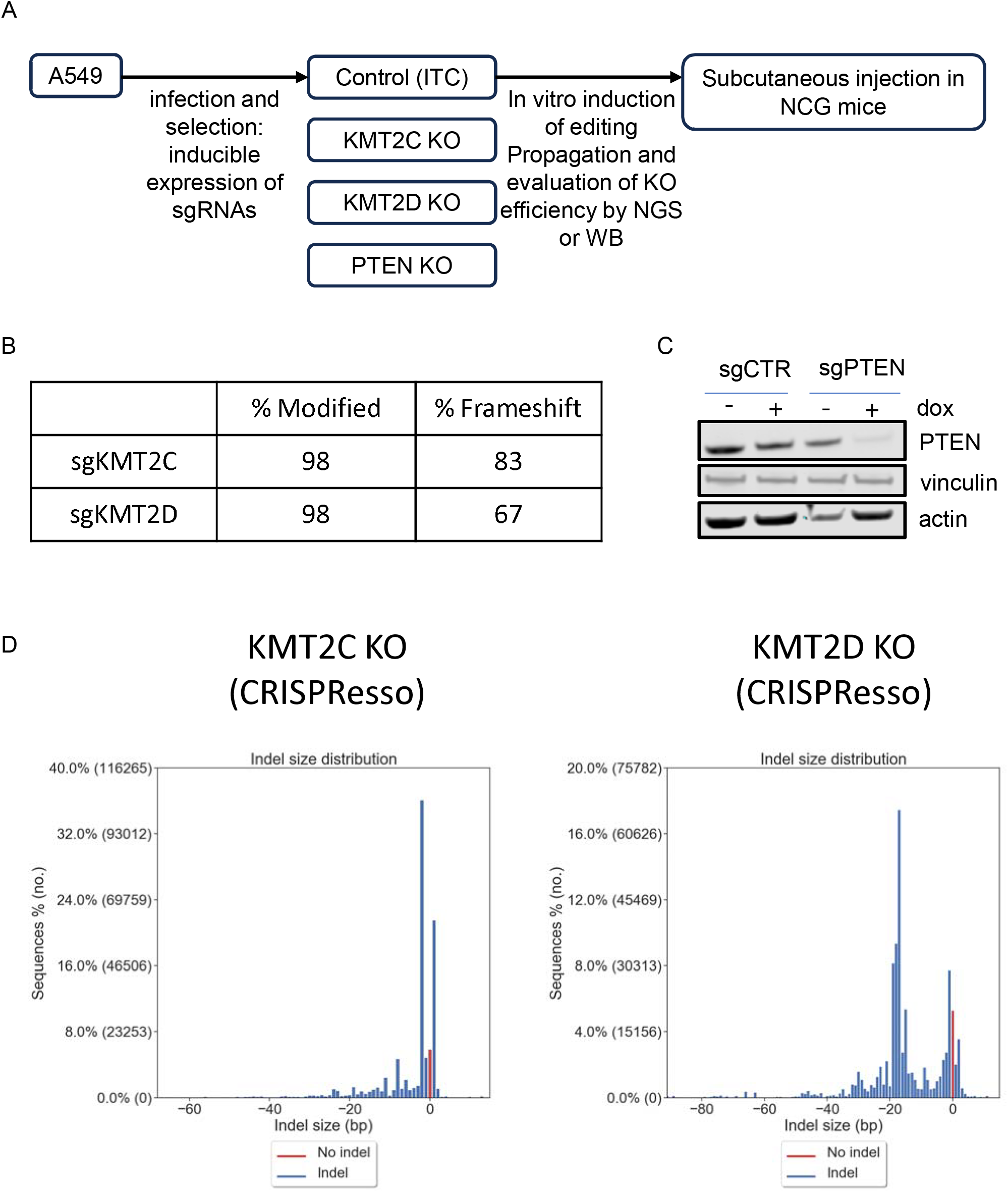
Editing efficiency analysis of KMT2C and KMT2D knockout in A549 cells. (A) Scheme of A549 isogenic cell line generation and *in vivo* validation experiment. (B) Editing efficiency analysis of KMT2C and KMT2D KO cells using CRISPResso^55^ analysis of. (C) Western blot validation of of PTEN knock out in isogenic lines. (D) Indel size distribution for KMT2C and KMT2D KO cells as determined by CRISPResso^55^

